# Neurons expressing mu opioid receptors of the habenula promote negative affect in a projection-specific manner

**DOI:** 10.1101/2021.09.13.460041

**Authors:** Julie Bailly, Florence Allain, Chloé Tirel, Florence Petit, Emmanuel Darcq, Brigitte Kieffer

## Abstract

**BACKGROUND:** The mu opioid receptor (MOR) is central to hedonic balance, and produces euphoria by engaging reward circuits. MOR signaling may also influence aversion centers, and notably the medial habenula (MHb) where the receptor is highly dense, however this was not investigated. Our prior data suggest that the inhibitory activity of MOR in the MHb limits aversive states. Here we therefore tested the hypothesis that neurons expressing MOR in the MHb (MHb-MOR neurons) promote negative affective states.

**METHODS:** Using *Oprm1*-Cre knock-in mice, we combined tracing and optogenetics with behavioral testing to investigate consequences of MHb-MOR neuron stimulation in approach/avoidance (real-time place preference), anxiety-related responses (open field, elevated plus maze and marble burying) and despair-like behavior (tail suspension).

**RESULTS:** Opto-stimulation of MHb-MOR neurons elicited avoidance behavior, demonstrating that these neurons promote aversive states. Anterograde tracing showed that, in addition to the interpeduncular nucleus (IPN), MHb-MOR neurons project to the dorsal raphe nucleus (DRN), uncovering a yet unreported connection of MHb to a main mood center. Opto-stimulation of MHb-MOR/IPN neurons triggered avoidance and despair-like responses with no anxiety-related effect, whereas light-activation of MHb-MOR/DRN neurons increased levels of anxiety with no effect on other behaviors, revealing two dissociable pathways controlling negative affect.

**CONCLUSIONS:** This study demonstrates aversive activity of MHb neurons that respond to MOR opioids. We propose that inhibition of these neurons by endogenous or exogenous opioids relieves negative affect via two distinct MHb microcircuits, contributing to despair-like behavior (MHb-MOR/IPN) and anxiety (MHb-MOR/DRN). This mechanism has implications for hedonic homeostasis and addiction.

## INTRODUCTION

The endogenous opioid system regulates hedonic homeostasis, and the mu opioid receptor (MOR) is an essential actor in this process. This receptor is well-known to mediate the strong euphorigenic effects and abuse potential of medicinal and abused opioid drugs drugs (1, 2). The MOR also contributes to rewarding effects of non-opioid drugs of abuse (3, 4), as well as social and food reward (5, 6), and pharmacological MOR blockade is aversive in both animals and humans, demonstrating that endogenous MOR signaling has rewarding activity (7, 8). Overall, it is well established that MOR activation either by opioid drugs or endogenously positively influences the hedonic tone.

Local pharmacological experiments have demonstrated that MOR activation is rewarding at several brain sites (9), and the positive hedonic effects of MOR activity have been particularly studied within the mesolimbic dopamine circuitry, considered a main reward pathway (10). There are, however, many other hedonic hot spots in the brain where MOR is expressed (9), therefore broader brain circuits are involved. Further, MOR may act at the level of both reward and aversion brain pathways (11) to either facilitate reward processing or limit the impact of aversive stimuli (8). The latter mechanism, however, has been poorly explored.

Intriguingly, MOR is densely expressed in the habenula (Hb) (12, 13), recognized as a major aversion center (14). This highly conserved brain structure is divided in a medial (MHb) and a lateral (LHb) subdivision (15) and the Hb complex connects forebrain to midbrain regions, to integrate cognitive with emotional and sensory processing (13, 16). The MOR is expressed predominantly in the MHb, composed of excitatory cholinergic and Substance P neurons projecting to the interpeduncular nucleus (IPN) (16). Several studies have suggested a role for the MHb in drug withdrawal, considered a strong aversive state. In particular, the MHb showed hyperactivity during nicotine withdrawal (17-19) and the nicotinic system in the MHb was shown to functionally interact with the opioid system in naltrexone-precipitated withdrawal (20-22). Thus, MOR signaling in the habenula likely influences the postulated aversive activity of MHb neurons.

We recently deleted the MOR gene in Chnrb4-positive neurons of the MHb, eliminating about half MORs in this brain structure, and showed that conditioned place aversion produced by naloxone administration was less well detected in mutant mice (23). This was a first genetic evidence that aversive effects of naloxone are mediated, at least in part, by MORs operating at the level of the MHb circuitry. Because MORs are inhibitory receptors, this finding led us to hypothesize that endogenous MOR signaling normally acts as a brake on MHb neurons expressing the receptor (thereafter referred as to MHb-MOR neurons) to limit aversive states.

To clarify the mechanisms underlying this observation and test the hypothesis, we focused our attention on the MOR-related circuitry in the habenula. We used our newly created *Oprm1*-Cre mice (24) to manipulate MHb-MOR neurons and establish whether these neurons, which normally respond and are inhibited by MOR opioids, indeed encode a negative emotional state. Data from the present study first confirm the original observation, and then demonstrate that opto-activation of MHb-MOR neurons induces aversive states. Nature of the negative affect differs depending on whether these neurons project to the canonical IPN projection site, or to the dorsal raphe nucleus (DRN), which we identified as a novel MHb projection site.

## METHODS AND MATERIALS

### Animals

Both the conditional *Chnrb4*-MOR mouse line (23) and the *Oprm1*-Cre knock-in mouse line (24) were group housed (maximum of five mice per cage) in a temperature- and humidity-controlled animal facility (21 ± 2°C, 45 ± 5% humidity) on a 12 h dark/light cycle with food and water available *ad libitum*. All experiments were performed in accordance with the Canadian Council of Animal Care and by the Animal Care Committees.

### Drugs

Naloxone (Sigma) was dissolved in 0.9% NaCl and injected subcutaneously in a volume of 10 ml/kg.

### Stereotaxic surgery

Animals were anesthetized with 5% isoflurane for 5 min and maintained at 2% isoflurane. For optogenetic and electrophysiology experiments, adult *Oprm1*^Cre/Cre^ male mice were injected unilaterally with 300 nL of AAV2.EF1a.DIO.ChR2-mCherry (RRID:Addgene_20297) or AAV2.EF1a.DIO.mCherry in the habenula (AP: -1.35, ML: -1.55, DV -2.9/2.8/2.7; volume of injection: 3x 100 nL, angle 32°) and implanted above the habenula (AP: -1.35, DV: -2.5 from dura, ML: -1.55, angle 32°) or the IPN (AP: -3.2, ML: -0.5, DV: -4.7, angle 6°) or the DRN (AP: -4.15, DV: -3.3, ML: 0). The implant was secured using a first layer of Metabond followed by a layer of dental cement. Mice were allowed to recover for at least 5 weeks after infusion of virus before habituation to the optic cord and behavioral testing.

### Immunohistochemistry

Mice were light-stimulated for 3 minutes (at 473 nm, 10 mW, 20 Hz, 10 ms pulse width) in their home cage 90 minutes prior to perfusion to allow for c-fos expression. Mice were anesthetized with intraperitoneal injections of cocktail containing ketamine/xylazine/acepromazine. An intracardiac perfusion was performed with ∼ 10 ml of ice-cold 1× Invitrogen PBS (Thermo Fisher Scientific) pH 7.4 followed by ∼ 50 ml ice-cold 4% paraformaldehyde (PFA; Electron Microscopy Sciences) using a peristaltic pump at 10 ml/min. Brains were dissected and postfixed after 24 h at 4°C in the 4% PFA solution, cryoprotected at 4°C in 30% sucrose (Thermo Fisher Scientific) for 48 h, embedded in OCT compound (Thermo Fisher Scientific), frozen, and finally stored at −80°C. Brains were sliced into 30 μm coronal and sagittal sections using a cryostat (Leica), and sections were stored at 4°C in PBS. Immunohistochemistry was performed by washing the sections for 3 × 10 min in PBS, then for 3 × 10 min with PBS/Triton X-100 0.1% (PBS-T; Sigma-Aldrich), followed by 1 h in a blocking buffer (PBS, 3% normal donkey serum, Triton X-100 0.2%), each at room temperature with gentle agitation. Sections were incubated overnight in blocking buffer at 4°C with the following primary antibodies: 1:2000 c-fos (Cell signaling technology), 1:2000 ds-red (Takara Bio). Sections were then washed 3 × 10 min in PBS-T, incubated for 2 h at room temperature with appropriate Alexa Fluor-conjugated secondary antibodies. Sections were washed 3 × 10 min in PBS-T with gentle agitation, placed in PBS, and mounted on to glass slides with Mowiol (PolyScience) and DAPI (0.5 μg/ml; Thermo Fisher Scientific; RRID:<u>AB 2307445</u>). Consecutive tissue sections were used for counting c-fos^+^ cells respectively in habenula (4 sections) and IPN nucleus (3 sections). Images were acquired using an Olympus FV1200 confocal microscope. For each animal (n=5/groups for figure 2 and figure 4), equivalent sections were chosen for cell counting using Allen Brain Atlas as a reference. c-fos^+^ cells were counted assisted by Fiji Cell Counter on ImageJ and fold changes compared to control were calculated. From a total of 108, 43 mice were excluded from analysis due to incorrect virus injection or cannula placement.

### Image acquisition

For fluorescence microscopy, an Olympus IX73 microscope with 10× or oil-immersion 60× objective was used. For confocal microscopy imaging, an Olympus FV1200 microscope with 20× objective, was used to take z-stack images. Slides were scanned on the Olympus VS120 microscope with a 10× objective.

### Behavior

#### Conditioned place aversion to naloxone in Chnrb4-MOR mice

On test day 1 (pre-test), mice had free access for 15 minutes to two compartments different by their color, floor texture and spatial configuration. The compartment mice preferred on day 1 was next associated with naloxone (biased configuration). On test days 2, 3 and 4, mice were injected subcutaneously with saline in the morning and confined to the saline-paired compartment for 30 minutes. Mice were next injected subcutaneously with naloxone (10 mg/kg, 1 mg/ml) or saline in the afternoon and confined to the naloxone-paired compartment for 30 minutes. On test days 5, 8, 11, 16, 24, 30, 34, 37 and 73 (post-tests) mice had again free access to both compartments for 15 min. Time spent in the drug-paired compartment was compared between the pre-test and post-tests and served as a measure of aversion induced by naloxone. Of note, on days 24, 30, 34 and 37 only naloxone-treated mice were tested to evaluate the extinction of aversion to naloxone. This was not done in saline-treated mice as there was no CPA to extinguish. Then, in between these tests, previously naloxone-treated mice were exposed to an extinction paradigm which consisted of injecting mice with saline and to confine them again for 30 minutes AM and PM in alternate compartments. Mice received two extinctions sessions/day for 2 days before each test.

#### Optogenetic behavioral experiments in Oprm1-Cre mice

##### Timeline

Prior to behavioral testing, mice were habituated to handling for 3 consecutive days. The battery of behavioral testing was organized as follows: mice underwent real-time place testing for four consecutive days (day 1-4) and then performed one behavioral test every 3 days (1 day of testing followed by 2 days of rest) in the order described below:

##### Real time place preference

Mice were placed in a behavioral arena (black Plexiglas, 50 × 50 × 25 cm) divided into two identical chambers and allowed to explore freely the environment for 20 min. Using an Anymaze hardware controller connected to the laser, light stimulation was delivered when mice were in the light-paired chamber (light stimulation parameters: blue laser at 473 nm, 10 mW at 0, 5, 10 or 20 Hz (10 ms pulse width)). At the start of the session, the mouse was placed in the non-stimulated chamber. The time spent on the paired stimulation side was recorded via a CCD camera interfaced with the Anymaze (Stoelting) software.

##### Openfield

The open field apparatus was made of transparent plastic (41 × 41 cm) and divided into a central field (center, 25 × 25 cm) and an outer field (periphery). Individual mice were placed in the center of the open field at the start of the session. The open field test consisted of a 9-minute session divided in three alternating 3-minute epochs (OFF-ON-OFF epochs, with a blue laser at 473 nm, 10 mW, 20 Hz, 10 ms pulse width). The time spent in the center was recorded via a CCD camera interfaced with the Anymaze (Stoelting) software.

##### Marble burying test

Mice were placed in a transparent plastic box (40 cm×20 cm×30 cm, bedding depth: 3cm). The box contained 20 glass marbles (four rows of five marbles equidistant from one another). Mice were allowed to explore for 18-min with alternating 3-minute epochs (blue laser at 473 nm, 10 mW, 20 Hz, 10 ms pulse width). Mice started with either an ‘on’ or ‘off’ session in a counterbalanced manner. Mice were then carefully removed from the testing arena and the number of marbles buried was recorded. The marble burying index was arbitrarily defined as the following: 1 for marbles fully covered with bedding, 0.5 for marbles that were half buried and 0 for marbles not buried.

##### Elevated-plus-maze test

The elevated plus maze (Viewpoint) apparatus was made of grey plastic and consisted of a central platform (6 × 6 cm) with two open arms (37 × 6 cm) and two closed arms (37 × 6 × 30 cm). Proximal areas were defined as regions with distances from the center of the maze less than 15 cm. The maze was elevated 50 cm from the floor and illuminated at 40 lux. At the start of a session, mice were placed in the center of the central platform facing the open arm. Each session was divided into three alternating 3-minute epochs (OFF-ON-OFF epochs with a blue laser at 473 nm, 10 mW, 20 Hz, 10 ms pulse width). The time spent in the proximal open arms was recorded via a CCD camera interfaced with the Anymaze (Stoelting) software. Mice that fell off the maze were excluded from the analysis.

##### Tail suspension test

An automated tail-suspension apparatus (Bioseb, USA) was used to measure immobility. Mice were suspended by the tail, using non-irritating adhesive tape, to a hook connected to a strain gauge that transmitted the animal’s movements to a central unit that calculated the total duration of immobility during a 6-min test. Each session was divided into two alternating 3-minute epochs (OFF-ON epochs with a blue laser at 473 nm, 10 mW, 20 Hz, 10 ms pulse width). Mice climbing their tail during the testing period were excluded from analysis.

### Statistical analysis

All data are presented as the mean SEM. Statistical analysis was assessed using t tests or repeated-measures ANOVA. When ANOVA reached significance, a Bonferroni’s post hoc test was conducted. Significance was defined as *p < 0.05, **p < 0.01, and ***p < 0.001.

## RESULTS

### Aversion to naloxone is reduced when MORs of the habenula are deleted

Our previous results showed that conditional MOR deletion in the MHb reduces naloxone-induced conditioned place aversion (CPA) in both naive and morphine-dependent mice (23). We first consolidated the original finding in naïve mice, which is at the basis of our working hypothesis. Thus, we submitted the conditional *Chnrb4*-MOR mice (i. e. lacking MOR in cells expressing the β4 subunit of the nicotinic acetylcholine receptor present in the MHb only, and representing about 50% MHb-MOR neurons) and their control littermates to naloxone place conditioning, and tested both acquisition and extinction of the behavior (**Figure 1**).

**Figure 1:**
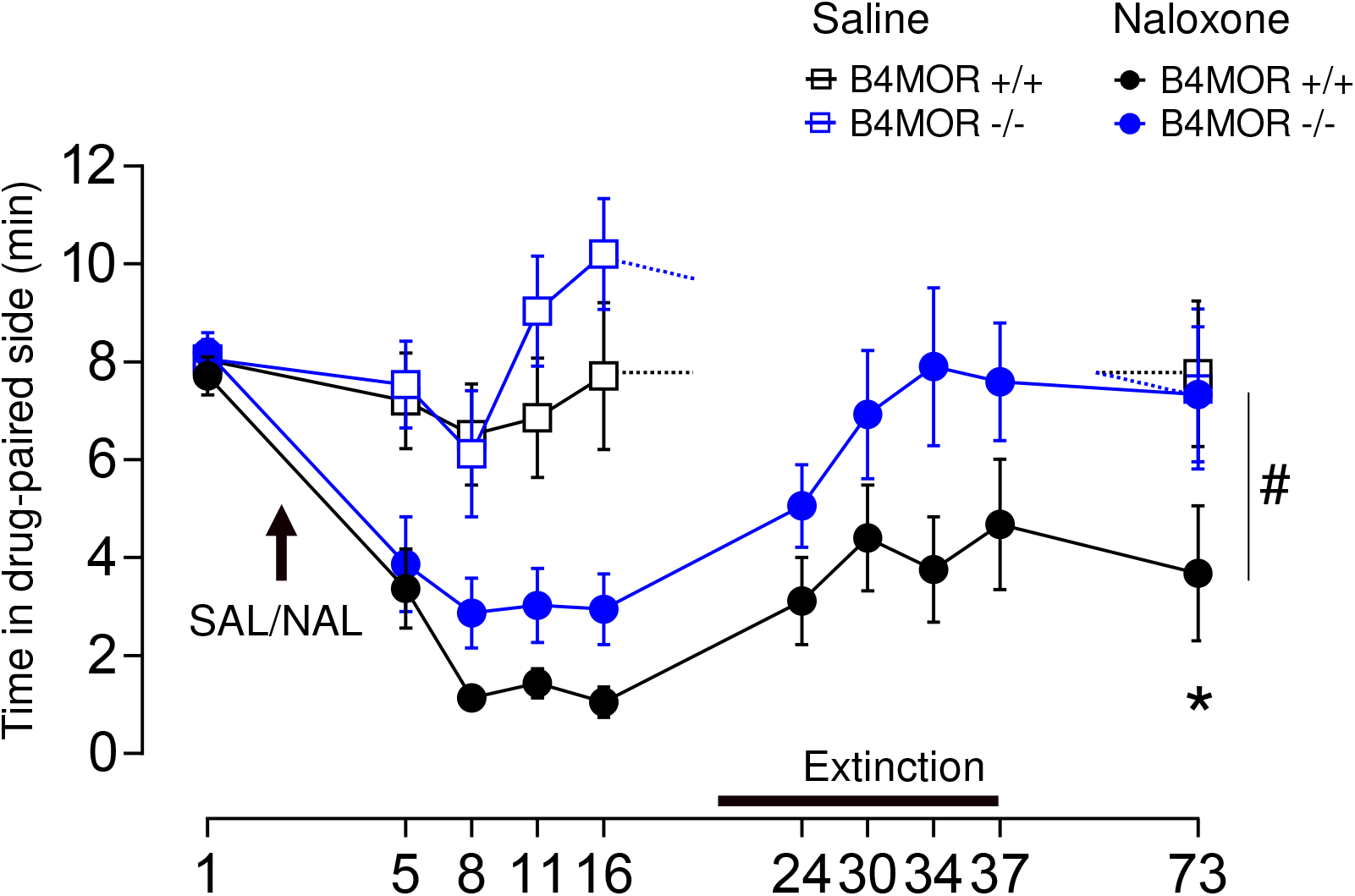
Naloxone-induced conditioned place aversion is reduced in Chnrb4-MOR mice. Chnrb4 mice and control littermates were submitted to place conditioning (SAL/NAL, days 2, 3 and 4). The two compartments were paired with either saline/saline (AM) or naloxone saline (PM) treatment (10 mg/kg). During testing (days 5, 8, 11, 16), mice freely explored the two compartments for 15 minutes. Two extinction sessions (saline/saline AM and PM) followed by testing were performed on days 24, 30, 34 and 37, and mice were finally tested one last time on day 73. Time spent in the drug-paired chamber is shown for the four groups. SAL, Saline. NAL, Naloxone 10-mg/kg/injection. N = 11-12/Group. #p < 0.01 Main effect of Genotype. *p < 0.05, test day 73 versus test day 1 in Chnrb4 mutant mice.

Naloxone (10 mg/kg) induced a conditioned place aversion (CPA) in both control and *Chnrb4*-MOR mice, which was detected along the entire testing procedure (three-way RM ANOVA on tests days 1-16 and 73; Main effect of Treatment, F_(1,43)_ = 29.09, \*\*\*\**p* < 0.0001) and this effect depended upon the test day (Treatment x Time interaction effect, F_(5,215)_ = 9.99, \*\*\*\**p* < 0.0001). Aversion to the naloxone-paired compartment was lower in *Chnrb4*-MOR mice compared to their Ctrl along the entire procedure (main effect of Genotype, F_(1,21)_ = 8.55, *p* = 0.008). We then extinguished the naloxone CPA during 4 sessions (days 24, 30, 34 and 37). Extinction of CPA to naloxone occurred in both genotypes and when a re-test was performed at day 73, only control mice still showed place aversion to naloxone (see statistical details in **Suppl Table S1**).

In conclusion, the data show acquisition of a long-lasting aversion for the naloxone-paired compartment in control animals, which was resistant to extinction. In contrast, *Chnrb4*-MOR mice showed a significant but lower aversion to the naloxone chamber, and this aversion did not persist after the extinction sessions. These data definitely confirm that the well-described aversive effect of naloxone results, at least partially, from blockade of endogenous MOR signaling in the MHb. Because MOR is an inhibitory receptor, MHb-MOR neurons should therefore promote aversive states and our next step was to test this hypothesis.

### Optogenetic stimulation of MHb-MOR neurons induces real-time place avoidance

We first tested the general behavioral effect of MHb-MOR neuron activation using optogenetics in a Real-Time Place Testing (RTPT) setting. We injected a Cre-dependent AAV2-channelrhodopsin virus in the MHb of *Oprm1*-Cre mice, and implanted an optic fiber above the MHb, in order to stimulate cell bodies of the entire MHb-MOR neuron population (**Figure 2A**). MHb-MOR/MHb-ChR2 mice spent significantly less time in the stimulation side at 20 Hz compared to the control group (unpaired t test, t_(19)_ = 2.484, *p<0.05 **Figure 2B-C**). The stimulation did not affect total activity measured at 20 Hz (unpaired t test, NS ; **Figure 2C**), indicating that no activity change could interfere with the avoidance response.

**Figure 2:**
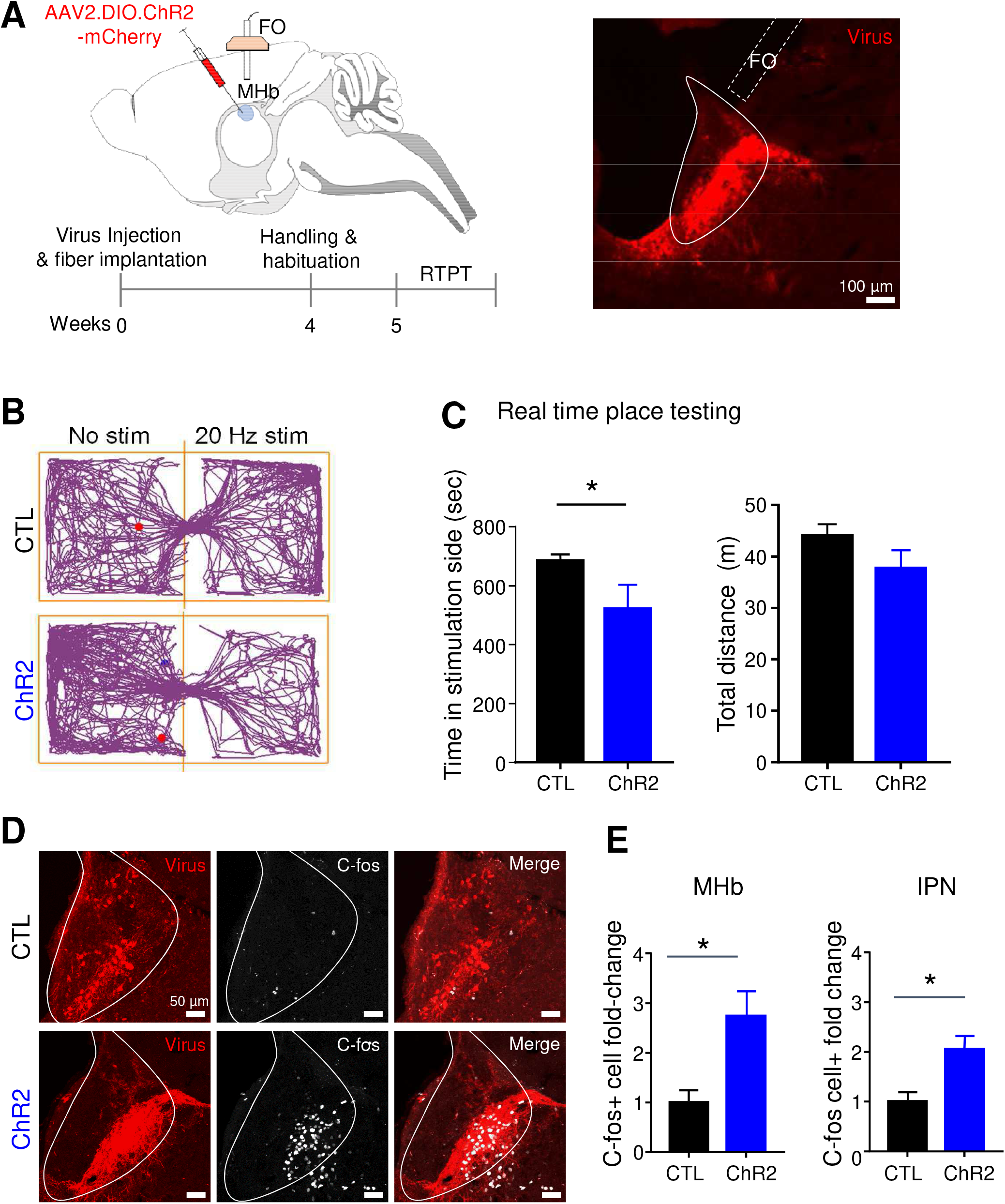
Opto-stimulation of MHb-MOR neurons induces avoidance behavior. *Oprm1*-Cre mice expressing either AAV2.EF1a.DIO.ChR2-mCherry (channelorhodopsin or ChR2) or AAV2.EF1a.DIO.mCherry (control or CTL) were subjected to real-time place preference testing. **A**. Scheme (left) and representative image (right) showing viral expression, with fiber-optic (FO) implantation above the MHb, and timeline for the experimental procedure. **B**. Representative tracking plot of the mice, at 20Hz (473 nm, 10 mW, 10 ms pulse width). **C**. Light stimulation in MHb-MOR/MHb-ChR2 mice induced significant behavioral avoidance to the light-paired side compared with control group (*n* = 6-12/group) without affecting the total distance traveled. Means are represented as ±SEM. *p<0.05. **D**. Confocal imaging of MHb sections demonstrating light-induced neuronal activation using c-fos as a marker of neuronal activity. ChR2-mCherry viral expression (red) and c-Fos immunostaining (white) are shown from representative sections. **E**. Quantification of c-Fos-positive cells upon light stimulation in the MHb and IPN of ChR2 and CTL animals.

To verify that the opto-stimulation conditions had successfully increased neuronal activity, we quantified c-fos expression using immunostaining (**Figure 2D-E**). We found a significant increase in the number of c-fos+ cells in response to a light stimulation in the brain of MHb-MOR/MHb-ChR2 mice compared to control animals, at the level of the implant (MHb, unpaired t-test, t_(9)_ = 3.007, *p > 0.01) and the main projection site (IPN, unpaired t-test, t_(9)_ = 3.218, *p < 0.01), confirming the efficacy of opto-stimulation parameters.

These data demonstrated that the light-activation of MHb-MOR neurons produces avoidance behavior in mice, suggestive of a negative emotional state.

### Hb-MOR neurons project to the IPN, and also to the DRN

We next verified whether MHb-MOR neurons project to the IPN, as postulated by the literature (16), and also tested whether other projection sites could be identified. We injected an anterograde Cre-dependent AAV2-mCherry virus in the MHb of *Oprm1*-Cre animals and amplified the signal by immunostaining after 4 weeks viral expression. As expected (12), we observed a strong fluorescent signal in both rostral and lateral parts of the IPN. Interestingly, we also identified significant fluorescence in the raphe nucleus, including dorsal raphe (DRN) and median raphe (MRN) nuclei (**Figure 3)**. This finding prompted us to investigate behavioral consequences of optogenetic stimulation for each MHb-MOR/IPN and MHb/-MOR/DRN projection.

**Figure 3:**
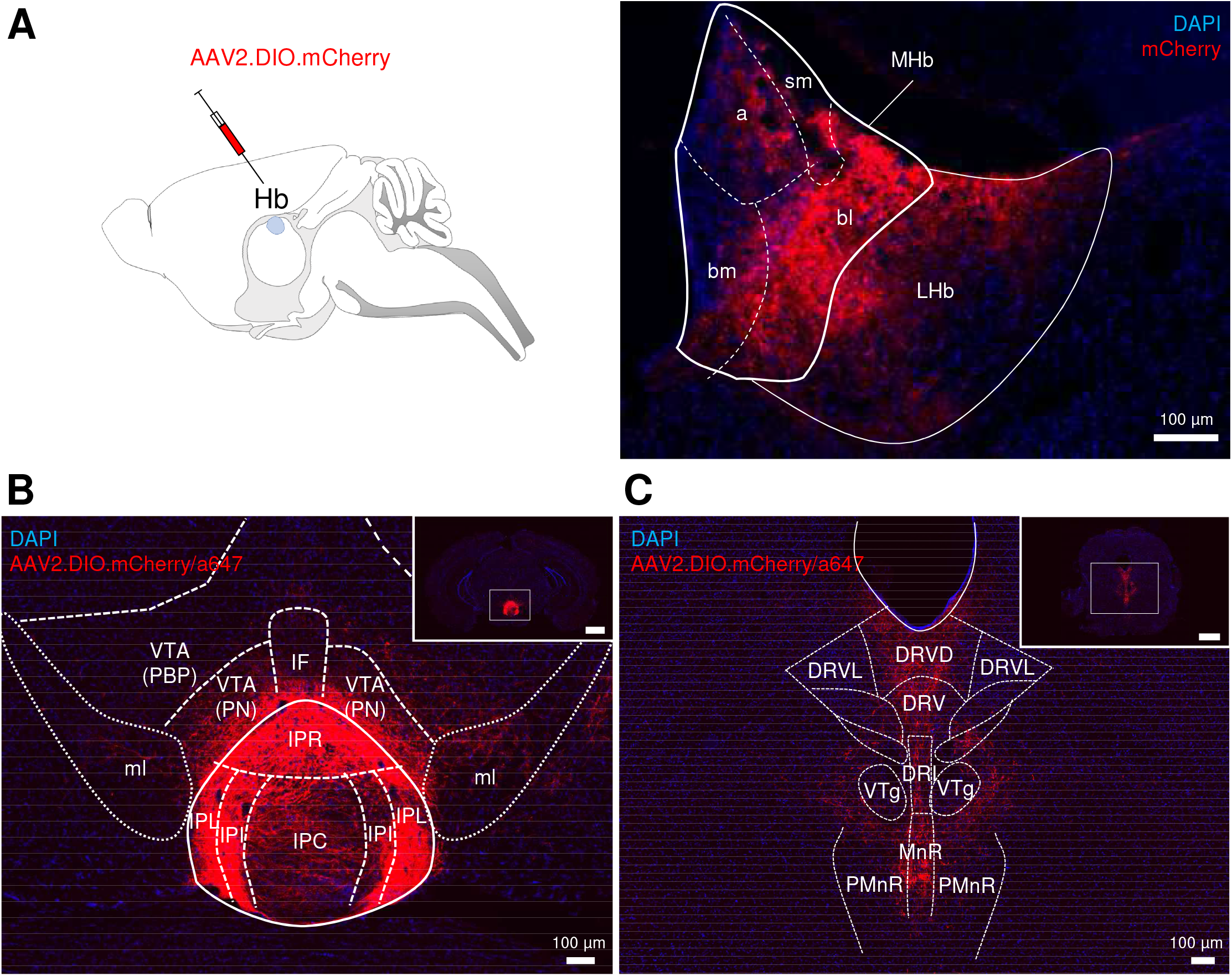
Identification of a novel projection site for MHb-MOR neurons. **A**. Anterograde tracing using a AAV2-DIO-mCherry virus was performed in the MHb of *Oprm1-Cre* mice, to visualize MHb-MOR projections (n=3). **B, C**. MHb-MOR neurons send most dense projections to the rostral and lateral part of the interpeduncular nucleus (B) as well as to raphe regions (C). a= apical, bl= basolateral, bm= basomedial, DRI/DRV/DRD/DRVL: dorsal raphe nucleus interfascicular, ventral, dorsal, ventral lateral, nucleus IPN/IPR/IPL/IPC= interpeduncular nucleus, rostral, lateral, intermediate, Hb= habenula, LHb= lateral habenula, MHb= medial habenula, ml= medial lemniscus, MnR= median raphe nucleus, PBP= parabrachial pigmented area of VTA, PMnR= paramedian raphe nucleus, PN= paranigral nucleus of VTA, sm= stria medullaris, VTA = ventral tegmental area, VTg= ventral tegmental nucleus.

### Optogenetic stimulation of MHb-MOR neurons to the IPN produces real-time place avoidance and a despair-like behavior

To test the role of MHb-MOR projecting to the IPN (MHb-MOR/IPN neurons), we injected the Cre-dependent AAV2 channelrhodopsin virus in the MHb of *Oprm1*-Cre mice and implanted an optic fiber above the IPN (**Figure 4A)**. Light-stimulation produced a strong place avoidance in the RTPT (**Figure 4 B-C**) (two-way RM ANOVA; significant frequency x virus interaction F_(3,66)_ = 7.468, ***p < 0.001). MHb-MOR/IPN-ChR2 mice spent significantly less time in the light-paired chamber at 10 Hz (**p < 0.01) and 20 Hz (***p < 0.001) compared to control mice indicating that optical stimulation was aversive. Again, the stimulation did not affect total activity measured at 20 Hz (unpaired t test, NS).

**Figure 4:**
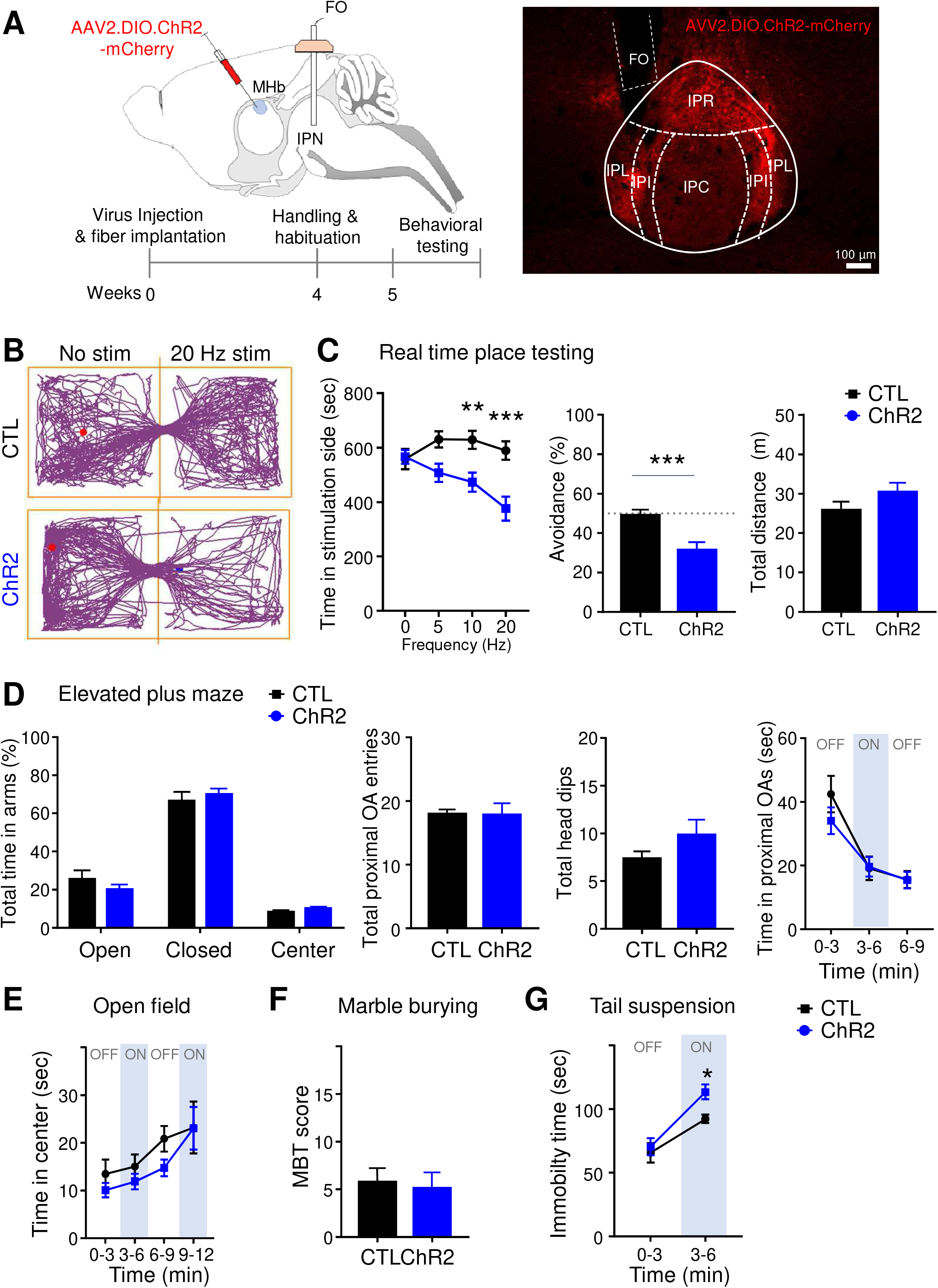
Optostimulation of MHb-MOR/IPN neurons induces avoidance and a despair-like behavior. *Oprm1*-Cre mice expressing either AAV2.EF1a.DIO.ChR2-mCherry (MHb-MOR/IPN-ChR2) or AAV2.EF1a.DIO.mCherry (CTL) were subjected to several behavioral tests to evaluate consequences of opto-activation on emotional responses. **A**. Scheme (left) and representative image (right) showing viral expression into the MHb, with fiber-optic (FO) implantation above the IPN, and timeline for the experimental procedure. **B**. Real-time place preference testing. Representative tracking plot of the mice at 20Hz (473 nm, 10 mW, 10 ms pulse width). **C**. (left) Light stimulation in MHb-MOR/IPN-ChR2 mice induced significant behavioral avoidance to the light-paired side compared with control group (n = 12/groups) and (right) the 20 Hz simulation condition triggered avoidance for the light-paired side without affecting the total distance traveled (NS). **D**. Elevated plus maze test. Activation of MHb-MOR/IPN neurons did not alter anxiety-level (n = 12/groups) for any of the tested parameters. Data are shown for the entire 9 min testing period, and see **Suppl Figure 1** for additional analysis. **E, F**. Open field and marble buying test revealed no effect of the light stimulation in MHb-MOR/IPN-ChR2 (n = 12/groups). **F**. Tail suspension test. Immobility time was significantly increased in MHb-MOR/IPN-ChR2 mice compared to control group (n=9-11/groups). Data are represented as mean values ±SEM. *p<0.05 **p < 0.01, ***p < 0.001.

In the elevated plus maze (**Figure 4D, Suppl Figure 1**), MOR-MHb/IPN-ChR2 mice spent the same amount of time in open arms, closed arms, and the center, and also did not differ from controls for other parameters (number of head dips in open arms, number of entries and time in proximal open arms). In the open-field (**Figure 4E)**, MHb-MOR/IPN-ChR2 mice and their controls spent the same amount of time in the center of the arena (two-way ANOVA, effect of virus, NS), and the two groups similarly increased time spent in the center as the test progressed and became less anxiogenic (two-way ANOVA, effect of time, F_(3,66)_ = 7.109, ***p<0.001). In the marble burying test (**Figure 4F)**, behavior did not differ between the two groups (unpaired t test, NS). Opto-stimulation of the MHb-MOR/IPN pathway therefore had no effect in any of the three anxiety-related tests.

In the tail suspension test (**Figure 4G)**, opto-stimulation significantly increased the mean immobility time for MHb-MOR/IPN-ChR2 mice compared to controls (two-way ANOVA, effect of virus, *p<0.05), suggestive of a despair-like behavior.

We finally verified again whether our conditions of optogenetic stimulation changes neuronal activity (**Suppl Figure 2A**). Light-stimulation of MHb-MOR/IPN significantly increased neuronal activity in the IPN (unpaired t test, t_(8)_ = 4.926, **p < 0.01), while no modification of c-fos staining was observed at the level of the MHb (unpaired t test, t_(8)_ = 0.8992, p > 0.05) (**Suppl Figure 2B**).This result indicated that opto-activation indeed modified activity of MHb-MOR neurons, which was detectable at the IPN terminals.

Altogether, these results demonstrate that selective stimulation of MHb-MOR neurons projecting to the IPN induces avoidance and despair-like behavior without altering levels of anxiety.

### Optogenetic activation of habenular MHb-MOR neurons to the DRN increases anxiety-related behavior

To test the role of MHb-MOR projecting to DRN (MHb-MOR/DRN neurons), we injected the Cre-dependent AAV2 channelrhodopsin virus in the DRN of *Oprm1*-Cre mice and implanted an optic fiber above the DRN (**Figure 5A)**.

**Figure 5:**
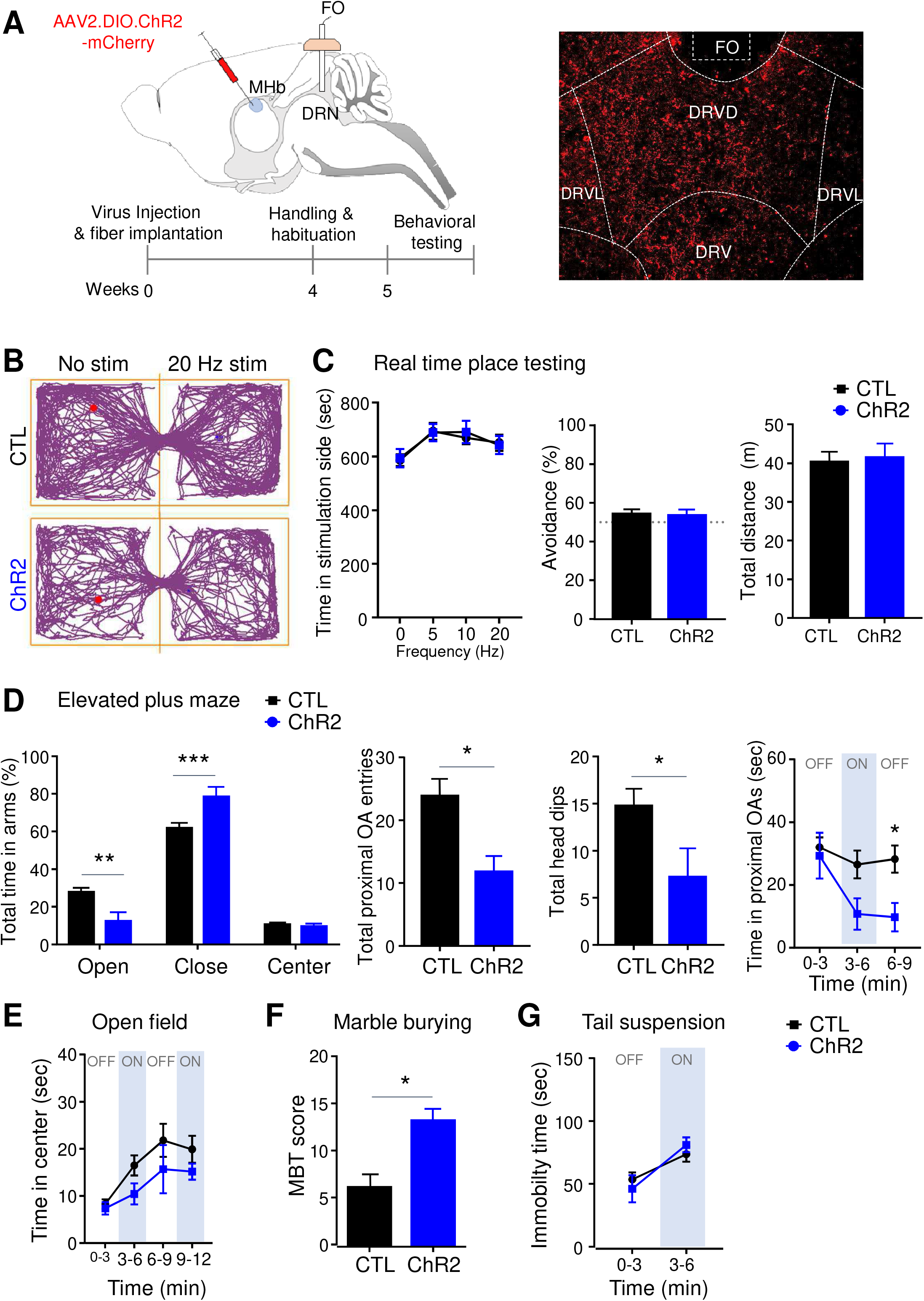
Optostimulation of MHb-MOR/DRN neurons increased anxiety-related behavior. *Oprm1*-Cre mice expressing either AAV2.EF1a.DIO.ChR2-mCherry (MHb-MOR/DRN-ChR2) or AAV2.EF1a.DIO.mCherry (CTL) were subjected several behavioral tests to evaluate consequences of opto-activation on emotional responses (as for MHb-MOR/IPN-ChR2 mice in **Figure 4). A**. Diagram (left) and representative image (right) showing viral delivery into the MHb and fiber-optic (FO) implantation above the DRN, a well as a timeline for the experimental procedure. **B**. Real time place preference testing. Representative tracking plot at 20Hz (473 nm, 10 mW, 10 ms pulse width). **C**. (left) Activation of MHb-MOR/DRN neurons does not produces place avoidance (n=6-14/groups) and (right) further analysis of the 20 Hz simulation condition indicated no alteration of the total distance traveled. **D**. Elevated plus-maze test. Activation of MHb-MOR/DRN neurons increases levels of anxiety for all the tested parameters (n=5-12/groups). Data are shown for the entire 9 min testing period, and see **Suppl Figure 1** for additional analysis. **E**. Openfield revealed no effect of the stimulation in MHb-MOR/DRN-ChR2 mice (n=6-14 /group). **F**. Marble burying test. Light-stimulation increased the burying (MBT) score in MHb-MOR/DRN-ChR2 (n=5-14 /group). **G**. Tail suspension test. There was no effect of the light-stimulation in MHb-MOR/DRN-ChR2 mice (n=5-14 /group). Data are represented as mean values ±SEM. *p<0.05 **p < 0.01, ***p < 0.001.

Optostimulation of this particular population of MHb-MOR neurons, under the same conditions as in previous experiments, did not trigger any significant effect in the RTPT for any group **(Figure 5 B-C**). Also there was no difference in time spent in the stimulation-side for MHb-MOR/DRN-ChR2 mice compared to controls at any frequency (two-way RM ANOVA, effect of group, NS). Finally, the stimulation did not affect total activity measured at 20 Hz (unpaired t, NS).

The optical stimulation, however, was anxiogenic. In the elevated plus maze test (**Figure 5D**), a two-way ANOVA revealed a significant interaction (F _(2,45)_ = 15.15, ***p < 0.001) for the % time in the two arms. MOR-MHb/IPN-ChR2 mice spent less time in open arms (**p<0.01) and more time in closed arms (***p<0.001) compared to control mice. In addition, the number of entries in proximal open arms and number of head dips were reduced in MHb-MOR/DRN-ChR2 compared to controls (unpaired t test, t_(15)_ = 2.613, *p<0.05; unpaired t test, t_(15)_ = 2.204, *p<0.05, respectively). Finally, the mean time spent in proximal open arms was lower for MHb-MOR/DRN-ChR2 mice during the 3-6 min stimulation period compared to control mice, a difference that became significant during the 6-9 min period of the test (*p<0.05) (see **Suppl Figure 3** for additional parameters). In the open-field (**Figure 5E**), MHb-MOR/DRN-ChR2 mice tended to spend less time in the center, although this effect did not reach statistical significance (two-way ANOVA, effect of groups, NS). Time spent in the center increased similarly for the two groups, as the test progressed and mice habituated to the open field (two-way ANOVA, effect of time, F_(3,54)_ = 6.745, ***p<0.001). In the marble burying test (Figure 5F), MHb-MOR/DRN-ChR2 mice displayed a higher marble index score compared to the control group (unpaired t test, t_(17)_ = 2.76, *p<0.01), suggesting an increased level of anxiety upon light-stimulation. The data together concur to demonstrate that opto-stimulation of the MHb-MOR/DRN pathway is anxiogenic, contrasting with data from the MHb-IPN neuron stimulation.

We examined effect of the opto-stimulation in the tail suspension test (**Figure 5G**). MHb-MOR/DRN-ChR2 mice and their controls showed the same time of immobility (two-way ANOVA, effect of groups, NS), suggesting that light stimulation of the DRN projection did not induce despair-like behavior, again contrasting with effects of the MHb-MOR/IPN neuron stimulation.

In sum, our data show that selective stimulation of MHb-MOR neurons projecting to the DRN has a strong anxiogenic effect, but does not seem to induce an aversive state that would result in place avoidance or despair-like behavior. Effects of opto-stimulation of this pathway therefore are the mirror image of those elicited by opto-stimulation of the MHb-MOR/IPN pathway, at least for emotional responses that we have tested.

## DISCUSSION

In sum (**Figure 6A**), we first confirm that naloxone aversion is reduced when the MOR is deleted from the MHb (23), setting the hypothesis that MOR, an inhibitory G protein coupled receptor, normally inhibits habenula function and thereby tempers its aversive activity. Second, we demonstrate that stimulation of habenular neurons expressing the MOR is aversive, substantiating the notion that endogenous MOR signaling in the MHb is anti-aversive. Third we show that MHb-MOR neurons project to the DRN, in addition to the classical projection to the IPN, discovering a new direct projection of MHb neurons to a main mood control center. Fourth, we demonstrate that MHb-MOR neurons projecting to either IPN or DRN modulate distinct emotional responses, uncovering two pathways controlling negative affect under the control of MOR opioids.

**Figure 6:**
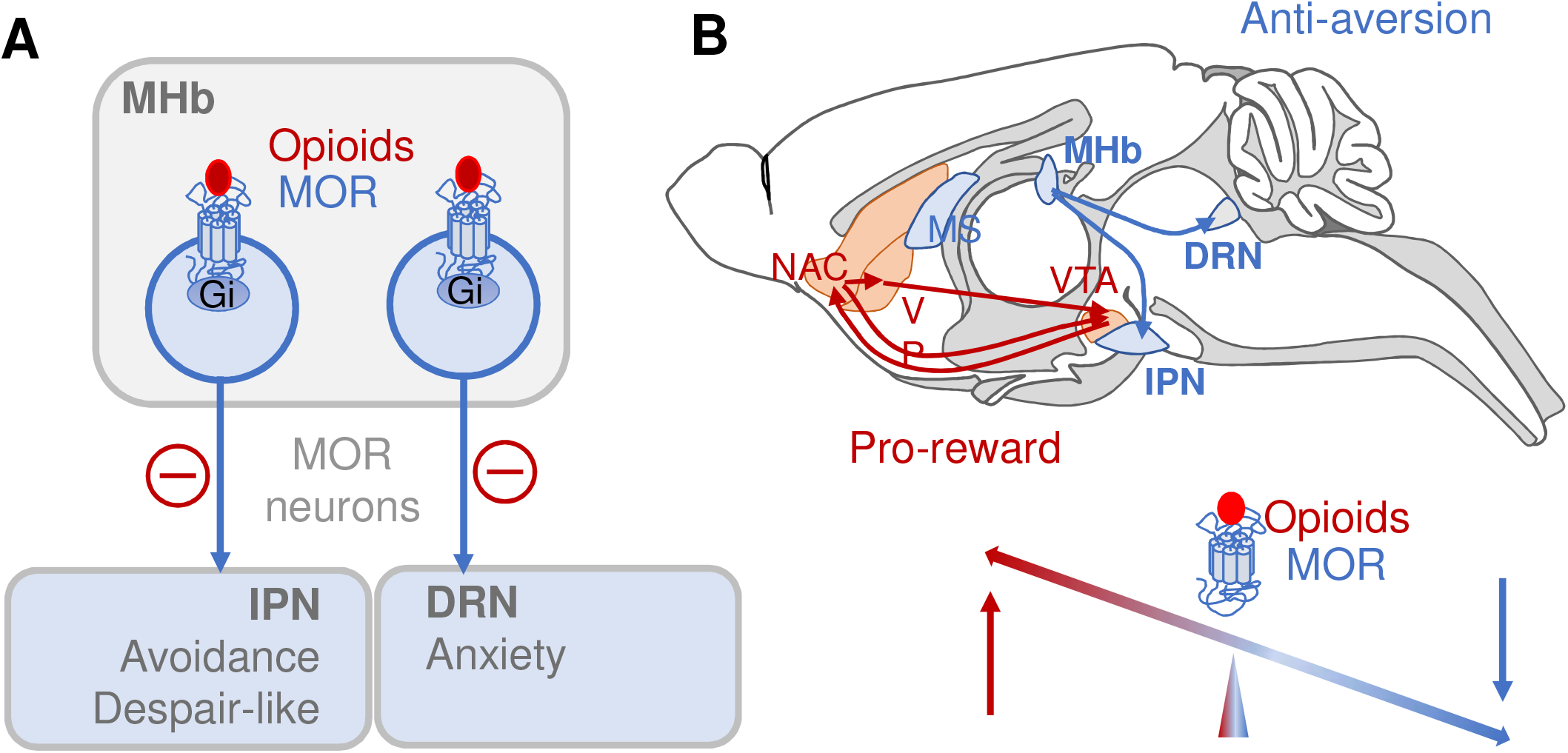
An anti-aversion mechanism for MOR signaling. **A**. This study demonstrates that (i) blockade of MORs in the MHb is aversive, (ii) stimulation of neurons expressing MOR (MHb-MOR) is aversive and (iii) selective stimulation of MHb-MOR/IPN and MHb-MOR/DRN pathways produces avoidance and despair-like behavior for the former, and enhances anxiety for the latter. We propose that endogenous or exogenous MOR signaling in the MHb, which inhibits MHb-MOR neurons (red negative sign), alleviates several aspects of negative affective states via at least two distinct circuits recruiting either the IPN or the DRN. MOR, mu opioid receptor; MOR neurons, neurons expressing the MOR; MHb, Medial Habenula; IPN, Interpeduncular Nucleus; DRN, Dorsal Raphe Nucleus. **B**. The pro-reward activity of MOR activation is well-established, and particularly well studied in the mesolimbic dopamine pathway (red). The present study reveals an “anti-aversion” function of MOR activity (blue) at the level of the MHb. Other pro-reward and anti-aversion MOR mechanisms exits in the brain, and together, these MOR-mediated mechanisms positively regulate hedonic homeostasis.

### MHb-MOR neurons encode an aversive state

MHb-MOR neurons represent about half of MHb neurons and form a unique ensemble of habenular neurons that respond to endogenous opioids and exogenous opiate drugs. Activity of these neurons therefore has implications for hedonic homeostasis, addiction and mood disorders. While the habenular complex is known critical in the emergence of aversive states (14), the specific role of the abundant MHb-MOR neurons has not been investigated earlier.

Here, we demonstrate that the activation of MHb-MOR cell bodies, i. e. the entire population of MHb-MOR neurons, produces an avoidance behavior (**Figure 2**). This is consistent with other findings demonstrating that MHb activity promotes negative affective states. Early studies showed that habenular lesions induce deficits in avoidance behavior induced by foot-shock (25).The overexpression of β4 receptor, enhancing MHb activity, resulted in a strong aversion to nicotine (26). On the contrary the knockout of several genes, known to contribute to habenular activity, reduce conditioned place aversion: deletion of CB1 receptors in the MHb inhibited expression of aversive memories (27), and knock-down of habenular RSK2, a protein contributing to intracellular signaling in MHb neurons (28) or knockout of GluN3A, a MHb-specific subunit of the excitatory NMDA receptor (29), both decreased lithium-induced conditioned place aversion.

Questions remain. The MHb is heterogeneous, composed of two main neuronal populations, substance P and cholinergic neurons, with regionally distinct distribution and possibly distinct roles (30). MOR-positive neurons overlap the two neuronal populations (12, 23), and whether these subpopulations play distinguishable roles in the aversive responses observed in the present study remains to be clarified. Also, our study demonstrates the existence of an anti-aversion opioid tone in the MHb (the naloxone experiment), as well as the negative affective activity of neurons expressing the MOR, however the origin and exact nature of endogenous opioid peptides acting at the MOR in this pathway, or the possibility of constitutive MOR activity at this brain site (31, 32), will need to be investigated in future studies.

### MHb-MOR neurons project to the raphe nucleus

While LHb connectivity is under intense investigation and expanding (33), less is known about the medial division. Our anterograde tracing experiments (**Figure 3**) showed that MHb-MOR neurons project to the rostral and lateral part of the interpeduncular nucleus (IPN), consistent with IPN being the main projection site for MHb (14), and our previous finding of high MOR expression in these IPN subdivisions (12). Interestingly we also found that MHb-MOR neurons project to several raphe structures including the DRN, which is home for serotonergic neurons and was also shown involved in morphine withdrawal (34, 35). In addition, anterograde fluorescence was observed in the median raphe nucleus, also containing serotoninergic neurons and involved memory consolidation, in interaction with the hippocampus (36). Overall, the finding of direct MHb connectivity to raphe nuclei is relatively new and consistent with data from the Allen Brain Atlas connectivity database (<u>Experiment 300236056</u>).

This MHb-raphe connection is of particular interest, as serotonin neurotransmission is key to mood control, negative affect and depressive behaviors (37, 38). The possibility that MHb neurons, and in particular those expressing MOR and responding to opioid stimulation, could directly influence raphe nuclei functions has strong implications for Opioids Use Disorders (OUDs), as understanding the negative affect of opioid withdrawal is key for therapeutic strategies (39). Of note, future studies are needed to determine whether the raphe projections, which we have identified, are collateral branches that diverge from the main MHb-IPN pathway or a separate circuitry. Our finding of dissociable behavioral responses modulated by each projection (see below) would favor the latter possibility.

### MHb-MOR neurons define two dissociable circuits

Several reports have demonstrated that the manipulation of septal afferences to the MHb, or the MHb itself, influences a diversity of emotional responses, including fear, anxiety and depressive-like behaviors (40-44). How do downstream projection sites of the MHb contribute to these functions, however, is virtually unknown, and another main finding of this work is that MHb-MOR-neurons promote aversive states in a projection-specific manner (**Figure 4 and 5**). Our data show that opto-activation of MHb-MOR/IPN neurons produces avoidance and a despair-like behavior, whereas a similar manipulation of MHb-MOR/DRN neurons increases levels of anxiety without effect on the other tested behaviors, highlighting two distinct circuits.

Consistent with the MHb-MOR/IPN finding, the pathogenesis of depression was correlated with increased IPN metabolism (45-47) and a recent study using a model of chronic stress demonstrated that hyperactivity of the MHb-IPN pathway is linked to depressive behavior, which was reversed by MHb lesion (48). We therefore speculate that activity of MHb-MOR/IPN neurons may generate a negative affect reminiscent to depression-related states. Less is known about a functional connection between the MHb and DRN. Recent studies in zebrafish and rats observed that alteration of MHb activity modifies both serotoninergic immunoreactivity or gene expression levels in the DRN (49, 50). Our own findings suggest that activity of the MHb-MOR/DRN circuit essentially facilitates anxiety states, distinguishable from depressive-like states elicited by the other circuit. This interpretation, however, is applicable only to the subpopulation MHb-MOR neurons, which we targeted in this study. Also, a strict separation of despair-type and anxiety-related responses is unlikely, as these complex processes overlap and for example, the silencing of cholinergic habenular inputs to the IPN did not alter basal anxiety level in naïve mice but reduced the exacerbated-anxiety in mice undergoing nicotine withdrawal (17).

Altogether, findings from this study indicate that MHb neurons responding to MOR opioids modulate several aspects of negative affective states, subserved by at least two distinct microcircuits. Our anterograde tracing experiments also revealed projections of MHb-MOR neurons to the lateral hypothalamus (not shown), which may also contribute and deserve further investigations.

### Conclusion: pro-reward and anti-aversion MOR functions

MOR is broadly expressed throughout reward and mood circuits, which regulate all aspects of hedonic homeostasis and are involved in addiction (2). To date, MOR opioids have been mostly associated to reward processes, and the “pro-reward” function of MOR is well-established. This is the first report demonstrating that morphine-responsive output neurons from the habenular complex, recognized as an anti-reward center, promote negative emotional responses and, in addition, via dissociable downstream pathways. Endogenous opioids may therefore alleviate several features of aversive states through this “anti-aversion” MOR mechanism. In other words, opioids may not only be “pro-reward” within the mesolimbic dopaminergic circuitry, as classically described, but also “anti-aversion” in the habenula circuitry, and perhaps other circuits, to overall modulate the hedonic balance (**Figure 6B**).

## Supporting information

Bailly et al. Supplemental

## ACKNOWLEDGEMENTS AND DISCLOSURE

We thank the staff at the animal facility of the Neurophenotyping Center and the Molecular and Cellular Microscopy Platform of the Douglas Mental Health University Institute (Montréal, Canada) for microscope usage. This work was supported by the National Institute of Health (# DA005010 and # DA048796 to BLK), the Canada Fund for Innovation and the Canada Research Chairs to ED and BLK, and by the Wallonie-Bruxelles International (JB).

All authors critically reviewed the content and approved the final version before submission. All data needed to evaluate the conclusions in the article are present in the article and/or supplementary material. Additional data related to this article are available upon request from the corresponding author. The authors report no biomedical financial interests or potential conflicts of interest.

## Author Contributions

JB, ED and BLK designed the study; FP performed animal care and genotyping; JB, FA and CT acquired the data; JB, FA, ED and BLK performed the analysis, interpreted data and wrote the manuscript. All authors read and approved the submitted version.

## Notes

### Competing Interest Statement

The authors have declared no competing interest.

