## Supplementary material for "Neurons expressing mu opioid receptors of the habenula promote negative affect in a projection-specific manner": Bailly et al. Supplemental

**Supplementary Information**

**Supplementary Table S1.** Statistical analyses for behavioral experiments in main figures.

| Figure | Stat | Factors | F (DFn, DFd) or t, df | p |
|---|---|---|---|---|
| **Figure 1** Aversion all conditions (Days 1-16 and 73) | Three-way RM ANOVA | Interaction (Treatment x Time) | $F_{5, 215} = 9.99$ | $p < 0.0001$ |
| | | Time | $F_{5, 215} = 11.52$ | $p < 0.0001$ |
| | | Treatment | $F_{1, 43} = 29.1$ | $p < 0.0001$ |
| | | Interaction (Treatment x Time x Genotype) | $F_{5, 215} = 1.32$ | $p = 0.26$ |
| | | Genotype | $F_{1, 43} = 2.79$ | $p = 0.1$ |
| **Figure 1** Aversion in NAL-treated mice only (all days) | Two-way RM ANOVA | Interaction | $F_{9, 189} = 0.9359$ | $p = 0.4952$ |
| | | Time | $F_{9, 189} = 10.64$ | $p < 0.0001$ |
| | | Genotype | $F_{1, 21} = 8.55$ | $p = 0.0081$ |
| **Figure 1** Extinction of aversion in NAL-treated mice only (Days 16-73) | Two-way RM ANOVA | Interaction | $F_{5, 105} = 0.5$ | $p = 0.78$ |
| | | Time | $F_{5, 105} = 5.86$ | $p < 0.0001$ |
| | | Genotype | $F_{1, 21} = 6.4$ | $p = 0.02$ |
| **Figure 1** Aversion day 1 vs 73 in B4MOR+/+ | Two-way RM ANOVA | Interaction | $F_{1, 21} = 3.42$ | $p = 0.079$ |
| | | Test | $F_{1, 21} = 4.52$ | $p = 0.046$ |
| | | Treatment | $F_{1, 21} = 4.02$ | $p = 0.058$ |
| **Figure 1** Aversion day 1 vs 73 in B4MOR-/- | Two-way RM ANOVA | Interaction | $F_{1, 22} < 0.5$ | $p = 0.92$ |
| | | Test | $F_{1, 22} < 0.5$ | $p = 0.52$ |
| | | Treatment | $F_{1, 22} < 0.5$ | $p = 0.99$ |
| **Figure 2c** Time in stimulation side | Student's *t-test* | | t=2.484 df=19 | $p$ =0.0225 |
| **Figure 2c** Total distance | Student's *t-test* | | t=1.552 df=19 | $p$ =0.1372 |
| **Figure 2e** Cells+ c-fos MHb | Student's *t-test* | | t=3.007 df=9 | $p$ =0.0148 |
| **Figure 2e** Cells+ c-fos IPN | Student's *t-test* | | t=3.218 df=9 | $p$ =0.0105 |
| **Figure 4c** Time in stimulation side | Two-way RM ANOVA | Interaction | $F_{3,66} = 7.468$ | $p$ =0.0002 |
| | | Time | $F_{3,66}= 5.338$ | $p$ =0.0024 |
| | | Virus | $F_{1,22} = 9.854$ | $p$ =0.0048 |

| | | | | |
|---|---|---|---|---|
| **Figure 4c** Avoidance | Student's *t-test* | | t=3.823 df=22 | *p* =0.0009 |
| **Figure 4c** Total distance | Student's *t-test* | | t=1.445 df=22 | *p* =0.1627 |
| **Figure 4d** Total time in arms | Two-way ANOVA | Interaction | $F_{2,66}$ = 1.016 | *p* =0.3676 |
| | | Arms | $F_{2,66}$= 179.1 | *p* <0.0001 |
| | | Virus | $F_{1,66}$ = 0.0001 | *p* =0.9996 |
| **Figure 4d** Total proximal OA entries | Student's *t-test* | | t=0.0429 df=22 | *p* =0.9662 |
| **Figure 4d** Total head dips | Student's *t-test* | | t=1.428 df=22 | *p* =0.1672 |
| **Figure 4d** Time in proximal OA arms | Two-way RM ANOVA | Interaction | $F_{2,44}$ = 1.101 | *p* =0.3415 |
| | | Time | $F_{2,44}$ = 26.69 | *p* <0.0001 |
| | | Virus | $F_{1,22}$ = 0.5067 | *p* =0.4841 |
| **Figure 4e** Time in center | Two-way RM ANOVA | Interaction | $F_{3,66}$ = 0.4059 | *p* =0.7492 |
| | | Time | $F_{3,66}$ = 7.109 | *p* =0.0003 |
| | | Virus | $F_{1,22}$ = 1.129 | *p* =0.2996 |
| **Figure 4f** Marble burying score | Student's *t-test* | | t=0.2933 df=20 | *p* =0.7723 |
| **Figure 4g** Immobility time | Two-way ANOVA | Interaction | $F_{1,34}$ = 1.822 | *p* =0.1860 |
| | | Laser OFF/ON | $F_{1,34}$ = 33.26 | *p* <0.0001 |
| | | Virus | $F_{1,34}$ = 4.586 | *p* =0.0395 |
| **Figure 5c** Time in stimulation side | Two-way RM ANOVA | Interaction | $F_{3,54}$ = 0.1531 | *p* =0.9273 |
| | | Time | $F_{3,54}$ = 6.785 | *p* =0.0006 |
| | | Virus | $F_{1,18}$ = 0.0130 | *p* =0.9104 |
| **Figure 5c** Avoidance | Student's *t-test* | | t=0.1686 df=18 | *p* =0.8680 |
| **Figure 5c** Total distance | Student's *t-test* | | t=0.2451 df=18 | *p* =0.8091 |
| **Figure 5d** Total time in arms | Two-way ANOVA | Interaction | $F_{2,45}$ = 15.15 | *p* <0.0001 |
| | | Arms | $F_{2,45}$ = 238.7 | *p* <0.0001 |
| | | Virus | $F_{1,45}$ = 0.0001 | *p* =0.9916 |
| **Figure 5d** Total proximal OA entries | Student's *t-test* | | t=2.613 df=15 | *p* =0.0196 |
| **Figure 5d** Total head dips | Student's *t-test* | | t=2.204 df=15 | *p* =0.0435 |
| **Figure 5d** Time in proximal OA arms | Two-way RM ANOVA | Interaction | $F_{2,30}$ = 2.074 | *p* =0.1433 |
| | | Time | $F_{2,30}$ = 5.437 | *p* =0.0097 |
| | | Virus | $F_{1,15}$ = 5.19 | *p* =0.0378 |
| **Figure 5e** Time in center | Two-way RM ANOVA | Interaction | $F_{3,54}$ = 0.579 | *p* =0.6313 |

| | | | | |
|---|---|---|---|---|
| | | Time | $F_{3,54} = 6.745$ | *p* =0.0006 |
| | | Virus | $F_{1,18} = 2.746$ | *p* =0.1148 |
| **Figure 5f** Marble burying score | Student's *t-test* | | t=2.76 df=17 | *p* =0.0134 |
| **Figure 5i** Immobility time | Two-way ANOVA | Interaction | F (1,34) = 0.9311 | *p* =0.3414 |
| | | Laser OFF/ON | $F_{1,34} = 12.67$ | *p* =0.0011 |
| | | Virus | $F_{1,34} = 0.0037$ | *p* =0.9952 |

**Supplementary Table S2**. Statistical analyses for behavioral experiments in Suppl Figures.

| Figure | Stat | Factors | F (DFn, DFd) or t, df | p |
|---|---|---|---|---|
| **Figure S1a** Total time in arms | Two-way ANOVA | Interaction | $F_{2,66}$ = 1.016 | *p* =0.3676 |
| | | Arms | $F_{2,66}$ = 179.1 | *p* <0.0001 |
| | | Virus | $F1_{,66}$ = 0.0001 | *p* =0.9996 |
| **Figure S1a** # total entries in arms | Student's *t-test* | Open | t=0.1614 df=22 | *p* =0.8732 |
| | | Closed | t=1.825 df=22 | *p* =0.0816 |
| **Figure S1a** # total proximal OA entries | Student's *t-test* | | t=0.0429 df=22 | *p* =0.9662 |
| **Figure S1a** # total head dips | Student's *t-test* | | t=1.428 df=22 | *p* =0.1672 |
| **Figure S1a** Total distance | Student's *t-test* | | t=1.732 df=22 | *p* =0.0973 |
| **Figure S1b** Time in arms (%) | Two-way ANOVA | Interaction | $F_{2,66}$ = 0.4719 | *p* =0.6259 |
| | | Arms | $F_{2,66}$ = 121.1 | *p* <0.0001 |
| | | Virus | $F_{1,66}$ = 0 | *p* >0.9999 |
| **Figure S1b** # entries in arms | Student's *t-test* | Open | t=3.168 df=22 | *p* =0.0045 |
| | | Closed | t=2.986 df=22 | *p* =0.0068 |
| **Figure S1b** # proximal OA entries | Student's *t-test* | | t=2.3996 df=22 | *p* =0.0253 |
| **Figure S1b** # head dips | Student's *t-test* | | t=0.817 df=22 | *p* =0.4227 |
| **Figure S1b** Distance | Student's *t-test* | | t=2.23 df=22 | *p* =0.0363 |
| **Figure S2b** Cells+ c-fos MHb | Student's *t-test* | | t=0.8992 df=8 | *p* =0.3948 |
| **Figure S2b** Cells+ c-fos IPN | Student's *t-test* | | t=4.926 df=8 | *p* =0.0012 |
| **Figure S3a** Total time in arms | Two-way ANOVA | Interaction | $F_{2,45}$ = 15.15 | *p* <0.0001 |
| | | Arms | $F_{2,45}$ = 238.7 | *p* <0.0001 |
| | | Virus | $F_{1,45}$ = 0.0001 | *p* =0.9916 |
| **Figure S3a** # total entries in arms | Student's *t-test* | Open | t=2.252 df=15 | *p* =0.0398 |
| | | Closed | t=1.112 df=15 | *p* =0.2837 |
| **Figure S3a** # total proximal OA entries | Student's *t-test* | | t=2.613 df=15 | *p* =0.0196 |
| **Figure S3a** # total head dips | Student's *t-test* | | t=2.204 df=15 | *p* =0.0435 |
| **Figure S3a** | Student's *t-test* | | t=2.296 df=15 | *p* =0.0365 |

| | | | | |
|---|---|---|---|---|
| Total distance | | | | |
| **Figure S3b** Time in arms (%) | Two-way ANOVA | Interaction | $F_{2,45}$ = 20.11 | *p* <0.0001 |
| | | Arms | $F_{2,45}$ = 259.1 | *p* <0.0001 |
| | | Virus | $F_{1,45}$ = 0.0001 | *p* >0.9999 |
| **Figure S3b** # entries in arms | Student's *t-test* | Open | t=2.196 df=15 | *p* =0.0443 |
| | | Closed | t=0.4297 df=15 | *p* =0.6734 |
| **Figure S3b** # proximal OA entries | Student's *t-test* | | t=3.07 df=15 | *p* =0.0078 |
| **Figure S3b** # head dips | Student's *t-test* | | t=2.202 df=15 | *p* =0.0437 |
| **Figure S3b** Distance | Student's *t-test* | | t=1.704 df=15 | *p* =0.1090 |

**Supplementary Figures Legends**

**Supplementary figure 1: Activation of MHb-MOR/IPN neurons does not affect levels of anxiety in the elevated plus maze. A.** Analysis is shown for all the recorded parameters on the total session period (0 to 9 min), see **Figure 4**. Statistics show no effect of the light stimulation (**Suppl Table 2**). **B**. Additional analysis focused on the stimulation period (3 to 6 min). Statistics confirm that opto-stimulation did not alter the time spent in open and closed arms and the number of head dips. However, we noted that opto-stimulation had a detectable hyper-locomotor effect over this short time period, which likely impacted on the number of entries in all the arms (open, closed, proximal open) (n=12/groups). *p<0.05, **p<0.01.

**Supplementary figure 2: Activation of MHb-MOR/IPN neurons increased c-fos expression in the IPN**. **A.** Confocal imaging of MHb sections demonstrating light-induced neuronal activation using c-fos as a marker of neuronal activity. ChR2-mCherry viral expression (red) and c-Fos immunostaining (white) are shown from representative sections. **B.** Quantification of c-Fos-positive cells in the MHb and IPN, for ChR2 animals and controls (n=5/groups). **p<0,01.

**Supplementary figure 3: Activation of MHb-MOR/DRN neurons increases levels of anxiety in the elevated plus maze. A.** Analysis is shown for all the recorded parameters on the total session period (0 to 9 min), see **Figure 5**. Statistics show that opto-stimulation increases the time spent in open arms, decreases time spent in closed arms, decreases number of entries in open arms and proximal open arms, and decreases the number of head dips (**Suppl Table 2**). Of note, opto-stimulation reduced locomotion activity over this time period, an effect that was not significant over the 3-6 min time period. **B.** Additional analysis focused on the stimulation period (3-6 min). Statistics confirmed similar higher anxiety for all the parameters (n=6-14/groups). *p<0.05, **p<0.01, ***p<0.001, ****p<0.0001.

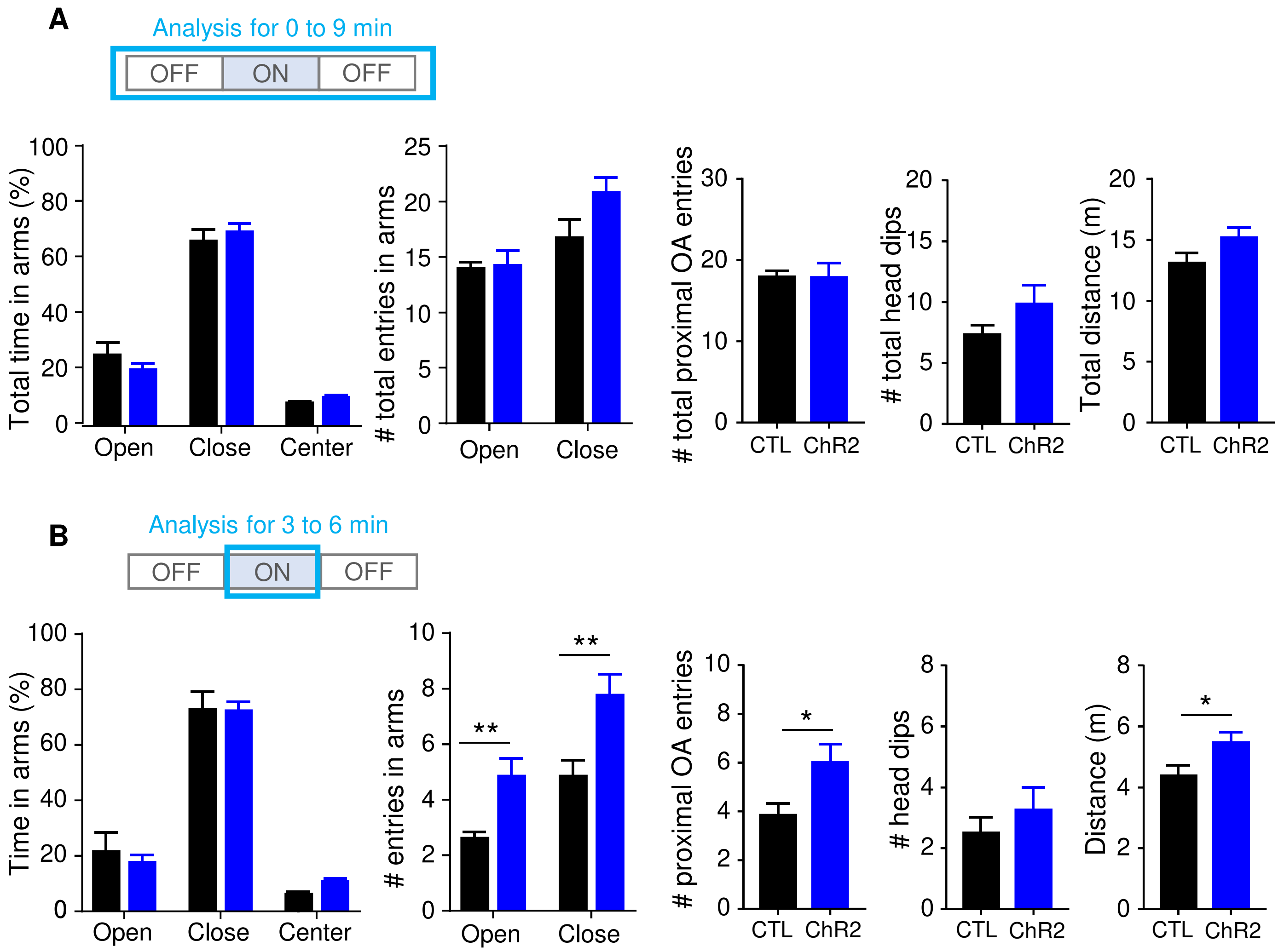

A
Analysis for 0 to 9 min
OFF
ON
OFF
Total time in arms (%)
Open
Close
Center
### total entries in arms
### total proximal OA entries
### total head dips
Total distance (m)
CTL
ChR2
B
Analysis for 3 to 6 min
Time in arms (%)
### entries in arms
### proximal OA entries
### head dips
Distance (m)
**
*

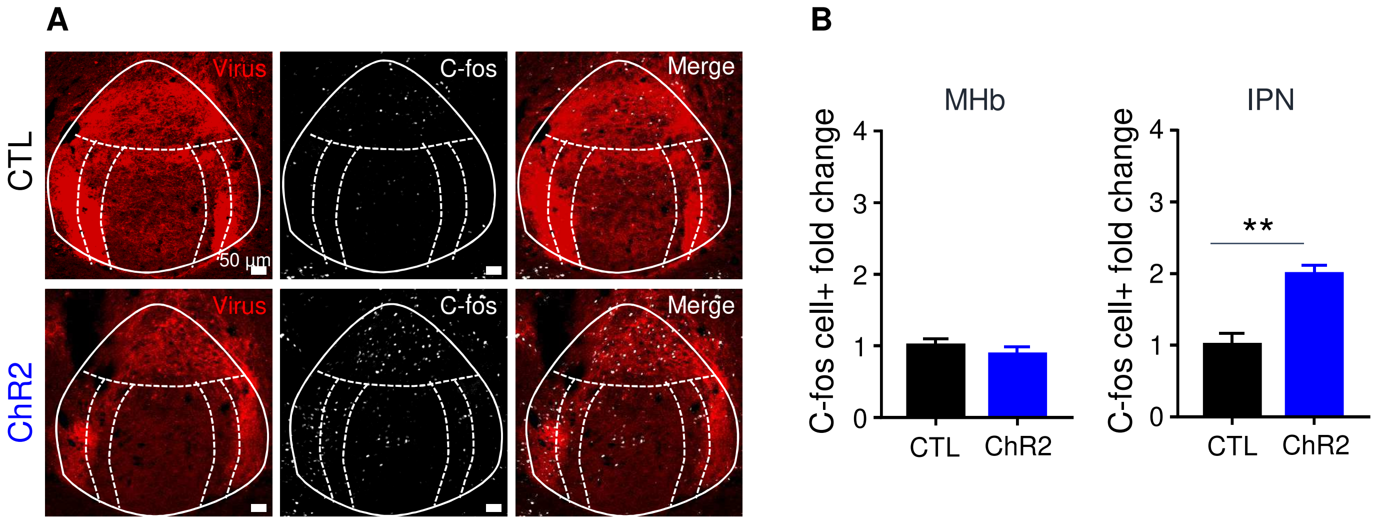
A
B
Virus
C-fos
Merge
CTL
50 μm
ChR2
MHb
IPN
C-fos cell+ fold change
4
3
2
1
0
**
CTL
ChR2

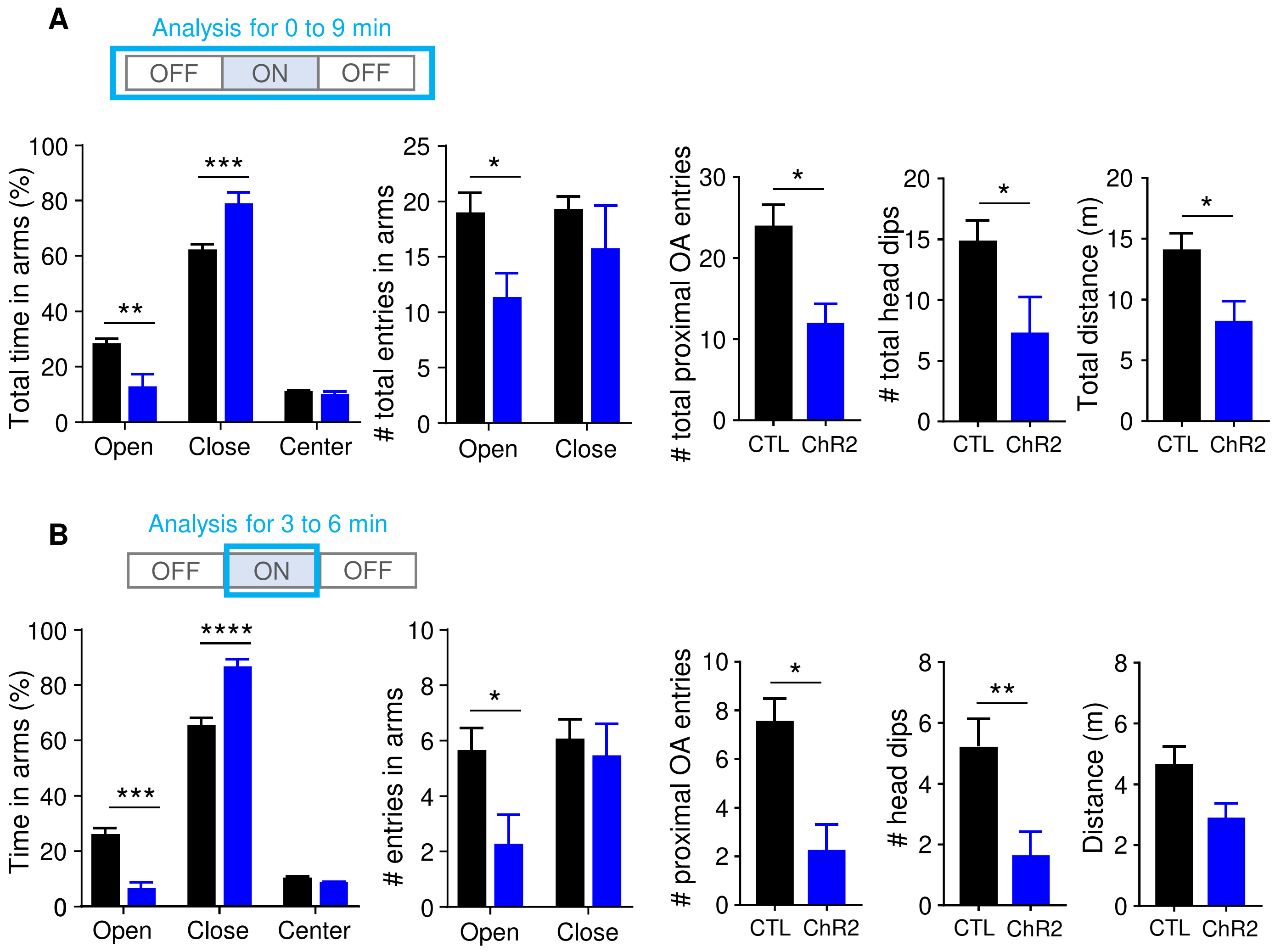

A
Analysis for 0 to 9 min
OFF
ON
OFF
Total time in arms (%)
100
80
60
40
20
0
**
***
Open
Close
Center
### total entries in arms
25
20
15
10
5
0
*
Open
Close
### total proximal OA entries
30
20
10
0
*
CTL
ChR2
### total head dips
20
15
10
5
0
*
CTL
ChR2
Total distance (m)
20
15
10
5
0
*
CTL
ChR2
B
Analysis for 3 to 6 min
OFF
ON
OFF
Time in arms (%)
100
80
60
40
20
0
***
****
Open
Close
Center
### entries in arms
10
8
6
4
2
0
*
Open
Close
### proximal OA entries
10
8
6
4
2
0
*
CTL
ChR2
### head dips
8
6
4
2
0
**
CTL
ChR2
Distance (m)
8
6
4
2
0
CTL
ChR2
